# Emergent Ecological Patterns and Modelling of Gut Microbiomes in Health and in Disease

**DOI:** 10.1101/2023.10.19.563037

**Authors:** Jacopo Pasqualini, Sonia Facchin, Andrea Rinaldo, Amos Maritan, Edoardo Vincenzo Savarino, Samir Suweis

## Abstract

Recent advancements in next-generation sequencing have revolutionized our understanding of the human microbiome. Despite this progress, challenges persist in comprehending the microbiome’s influence on disease, hindered by technical complexities in species classification, abundance estimation, and data compositionality. At the same time, recently the existence of macroecological laws describing the variation and diversity in microbial communities irrespective of their environment has been proposed using 16s data and explained by a simple phenomenological model of population dynamics. We here investigate the relationship between dysbiosis, i.e. in unhealthy individuals there are deviations from the “regular” composition of the gut microbial community, and the existence of macro-ecological emergent law in microbial communities. We first quantitatively reconstruct these patterns at the species level using shotgun data, offering a more biologically interpretable approach, and addressing the consequences of sampling effects and false positives on ecological patterns. We then ask if such patterns can discriminate between healthy and unhealthy cohorts.

Concomitantly, we evaluate the efficacy of different population models, which incorporate sampling and different ecological and statistical principles (e.g., the Taylor’s law and environmental noise) to describe such patterns. A critical aspect of our analysis is understanding the relationship between model parameters, which have clear ecological interpretations, and the state of the gut microbiome, thereby enabling the generation of synthetic compositional data that distinctively represent healthy and unhealthy individuals. Our approach, grounded in theoretical ecology and statistical physics, allows for a robust comparison of these models with empirical data, enhancing our understanding of the strengths and limitations of simple microbial models of population dynamics.

**Author summary:** In this study, we explore emerging ecological properties in gut microbiomes. Our aim here is to determine whether these patterns can be informative of the gut microbiome (healthy or diseased) and unveil essential ingredients driving its population dynamics. Leveraging on phenomenological models of species abundance fluctuations and metagenomics data, we highlight the pivotal role of Taylor’s law, a straightforward mathematical relation, in constructing theoretical models for the human gut microbiome. We thus explore such a general theoretical framework for investigating microbiome composition and show that not all ecological patterns are informative to characterize its states, while few are (e.g., species diversity). Eventually, thanks to the ecological interpretability of the inferred models’ parameters, our analysis provides insights into the role of environmental fluctuations and carrying capacities of the gut microbiomes in both health and disease. This study offers valuable knowledge, bridging theoretical concepts with practical implications for human health.

## Introduction

Next-generation sequencing has expanded our capacity to explore microbial biodiversity in a previously unachievable depth. This ’data explosion’ presents both challenges and exciting prospects. Over the past 15 years, biomedical researchers have leveraged this technology to delve into the human microbiome — the complex ecosystem of microorganisms coexisting in and on the human body [1–4]. This approach has illuminated countless microorganisms, once inaccessible via conventional culturing methods. All these efforts have aimed to establish a community resource program to build comprehensive reference datasets and develop computational tools and clinical protocols. Although several recent studies underscore the critical role of the microbiome in human health [5–11], our understanding of how the microbiome influences disease is still limited. Current methods, primarily focused on correlations and associations within the microbiome, are useful but often fail to identify the actual causes behind these patterns [12]. In part, this is also due to several technical challenges in species classification and abundance estimations, like sampling effects, false positives, and data compositionality. In fact, capturing only sample fragments of the entire genetic material leads to sparse datasets, where zeros abundances do not always imply species absence [13, 14]. Taxonomic profiling introduces false positives due to genome sequence overlaps, often resolved by selecting appropriate databases or setting abundance cut-offs [15]. Additionally, normalization, such as sum-to-one, is essential due to the compositional nature of microbiome data, significantly impacting data analysis [16–18]. Nevertheless, microbiomes data display several emergent ecological patterns that are suitable to be explained through population dynamics models. From probabilistic models like the Multinomial Dirichlet Distribution [19], which estimates relative abundances and considers sampling effects, to more complex interaction-based models like the generalized Lotka-Volterra model or other phenomenological/computational models based on inferring species interactions [20–23]. Eventually, non-interacting stochastic models reflecting basic ecological processes, like the stochastic logistic model [24], have demonstrated their effectiveness in reproducing microbiomes macroecological patterns [13].

However, an investigation of a possible link between such emergent patterns and dysbiosis is still missing. This study intends to address this gap by integrating theoretical frameworks in population dynamics modelling with empirical data on gut microbiomes in health and in disease. In particular, our work aims to: 1) Quantitatively reconstruct gut microbiome emergent patterns in health and in disease at the species level using shotgun data through a recently proposed taxonomic classifier [**?**] (previous studies used 16s data and OTU clustering [13], which are biologically less interpretable); 2) Examine the consequences of sampling effects and false positives on such ecological patterns; 3) Compare how well different models can describe such patterns and test possible differences between healthy and diseased cohorts; 4) Understand possible relationships of the inferred models parameters (having a well-defined ecological interpretation) with gut microbiome state (e.g., health or disease), so to be able to generate synthetic compositional data with statistically significant differences between healthy and unhealthy individuals.

In particular, our analysis includes a meta-analysis of studies on gut microbiomes in healthy individuals and those with gastrointestinal diseases [4, 10, 25], properly filtered. For both cases, we calculate the most relevant ecological patterns [26], and fit such patterns with different models. We consider three models. The Multinomial Dirichlet Distribution (MD) [19, 27], a statistical model including sampling effects and compositionality, but that does not satisfy Taylor’s Law [13, 28]; The Poisson stochastic logistic model (PSLM) [13, 24], incorporating sampling and fundamental processes (such as the Taylor law and environmental noise, but not compositionality. The Multinomial Symmetric Scaled Dirichlet Distribution (MSSD) [29, 30], which we show, combines both MD and PSLM through a general sampling theory. Our framework, grounded in theoretical ecology and statistical physics [31, 32], allows a robust comparison of these models with empirical data from studies on gut microbiomes in healthy individuals and those with gastrointestinal diseases, improving our understanding of the relative importance of the different mechanisms/processes considered, and also investigating the opportunity for such models to give insights on the gut dysbiosis.

## Materials and methods

### Theoretical Framework

We start by introducing the stochastic logistic model [13, 24], which gives the evolution of S species abundances in time

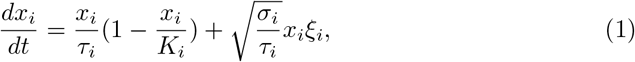

where *x*_*i*_ ∈ (0, *∞*), *K*_*i*_ is the carrying capacity of species *i* = 1 … *S, τ*_*i*_ sets the growth time scale and *σ*_*i*_ is the width of environmental noise experienced by the i-th species. The latter captures the fluctuations induced to the species growth rate by the environment (e.g. host) and by species interactions [33].

It can be shown that this process, once stationarity is reached, follows a Γ *Distribution*, i.e. 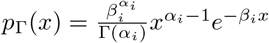, which describes the abundance fluctuations of species *i* among different samples, without considering compositionality [16, 34] and sampling effects [14]. Generally, we will refer to the distribution of abundance of a given species along different samples as abundance fluctuation distribution. The parameters *α*_*i*_ and *β*_*i*_ can be related to the mean and variance of the populations and the ecological parameters given by the model as

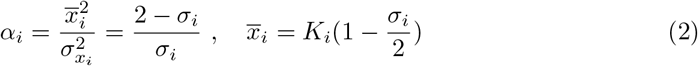

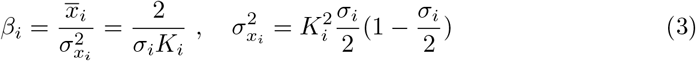

If we now consider compositionality and work with the relative populations 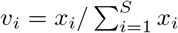, then the latter are distributed following the Scaled Dirichlet Distribution [29]

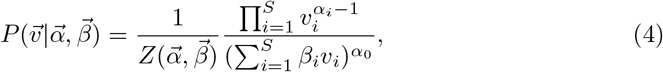

where the distribution is defined with the constraint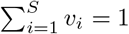. We have also introduced 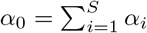, and the normalization constant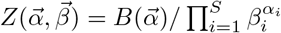.

Finally, 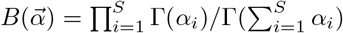 is the *S*-variate Beta function. An exhaustive derivation of this result can be found in Section 4 of the Supplementary Material. Regardless of the particular form 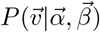 assumes, it describes the behavior of the *relative species abundances* in a given sample and lies in the (*S −* 1)-dimensional simplex Δ^*S*^. Such a family of distributions has 2S parameters. Since the number of observed species is high (*S* ∈ [10^2^, 10^3^]) and the microbiome dataset typically includes *R* ≈ 10^2^ samples, fitting this model is an unfeasible task. As we will show, we can greatly reduce the number of free parameters by constraining the model through relationships obtained from empirical macro-ecological patterns, as proposed by [13].

First, we consider the Taylor’s Law (TL). It takes the form of a scaling relation between the mean relative abundance of a species (among samples) and its fluctuations, i.e.

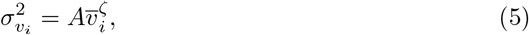

for *i* = 1 … *S*, where *A* and *ζ* do not depend on *i*. Compositionality does not affect this law if *ζ* = 2, and thus the same *A* is also found if we consider 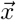, instead of 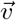 (see Supplementary Materials Section 4). Therefore, we can connect the ecological parameters *K*_*i*_, *σ*_*i*_ with *ζ* = 2, finding that TL is informative on both intra-species competition (driven by K) and the intensity of environmental noise (*σ*).

Exploiting Eqs. (2)-(3), we can thus reduce the number of parameters in our Scaled Dirichlet Distribution model to 2 + *S*. In fact, taking into account the parameterization of the Γ distribution in terms of *α* and *β*, the abundance fluctuation distribution depends only on its first moment 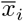, *ζ* and *A*:

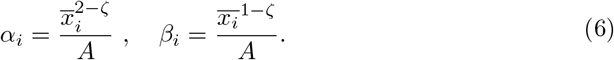

The dependence of the *α*s and *β*s on the exponent *ζ* suggests that there exist two interesting behaviors for the Scaled Dirichlet Distribution. In fact, for *ζ* = 1 we have a Poisson-like scaling as variance and mean are proportional, and the Scaled Dirichlet Distribution reduces to the Dirichlet distribution. The other limiting behavior with *ζ* = 2 is classically encountered in theoretical ecology [28]. In the following, we will only consider these two limiting cases, reducing the number of free parameters to 1 + *S*: one for TL’s amplitude *A*, while the remaining are for the mean abundances. From 6, we find that environmental fluctuations strength is the same for all species 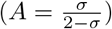 only in the case *ζ* = 2. Otherwise, the equation introduces a constraint for *K*_*i*_ and *σ*_*i*_, which would require a fine-tuning of the parameters to be satisfied. [13, 28]. Therefore, considering these values for *ζ*, after some manipulations, we obtain the following distributions:

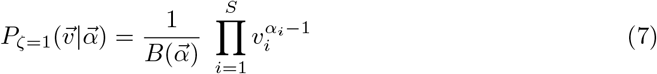

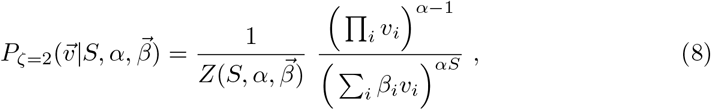

where 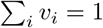. The *ζ* = 1 prescribes a Poisson-like scaling with 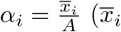 and *A* can be directly obtained from the data), and all the *β*s are proportional to an irrelevant constant. Finally, the corresponding distribution is the Dirichlet distribution. On the other hand, for *ζ* = 2 we find that all *α*_*i*_ = *α* = (2 *− σ*)/*σ* = *A*^*−*1^ are constant and 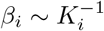. Due to the invariance of the corresponding distribution by rescaling of the *β*s, the proportionality constant is irrelevant. Also, because in the latter case *α*_*i*_ = *α*, then the dependence of 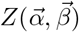 on 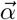 reduces to 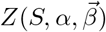 and we call the corresponding joint pdf the Symmetric Scaled Dirichlet distribution.

The second pattern that we exploit to reduce the number of parameters of the models is the species *mean* (relative) *abundance distribution* (MAD), which describes the frequencies of the average abundances. Indeed, it has been shown that different types of microbiomes share the same average (relative) species abundance distribution [13], i.e., 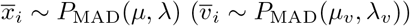 where the parameter *µ* (*µ*_*v*_) is defined so that 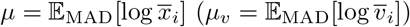; while λ (λ_*v*_) can be calculated analytically given the specific shape of *P*_MAD_, which solely depends on *µ* and λ. Typically, the distribution *P*_MAD_ has fat tails, and it is compatible with a Log-Normal distribution [13]. Actually, for the Symmetric Scaled Dirichlet distribution, we can show that the dependence on *µ* disappears, due to model symmetries (see Supplementary Materials, section 4). Regardless of the particular form *P*_MAD_ takes, its presence allows us to generate all the *S* averages once µ and λ are fitted from the data.

To take into account the effect of sampling, that in microbiome data cannot be neglected [13, 14], we introduce the convolution of the Dirichlet and Symmetric Scaled Dirichlet distribution with a multinomial one. In the first case, convolving the Dirichlet distribution with multinomial sampling, we obtain the Multinomial Dirichlet Distribution [19] (MD). The independent parameters of the model are the MAD parameters (*µ*,λ) and the TL amplitude *A*. In the second case, convolving the Symmetric Scaled Dirichlet distribution with the multinomial distribution, we obtain a novel model with ecologially grounded constraints, which we will refer to in the following as Multinomial Symmetric Scaled Dirichlet (MSSD). In this way, we can generate a synthetic microbiome using the relative abundance of species 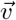 as the densities and the number of reads *N* as the number of trials of the multinomial distribution (see Fig. 1). In this case, the number of independent parameters to be fitted from the data is two, being λ and A (or, equivalently, λ and *σ*).

**Fig 1.**
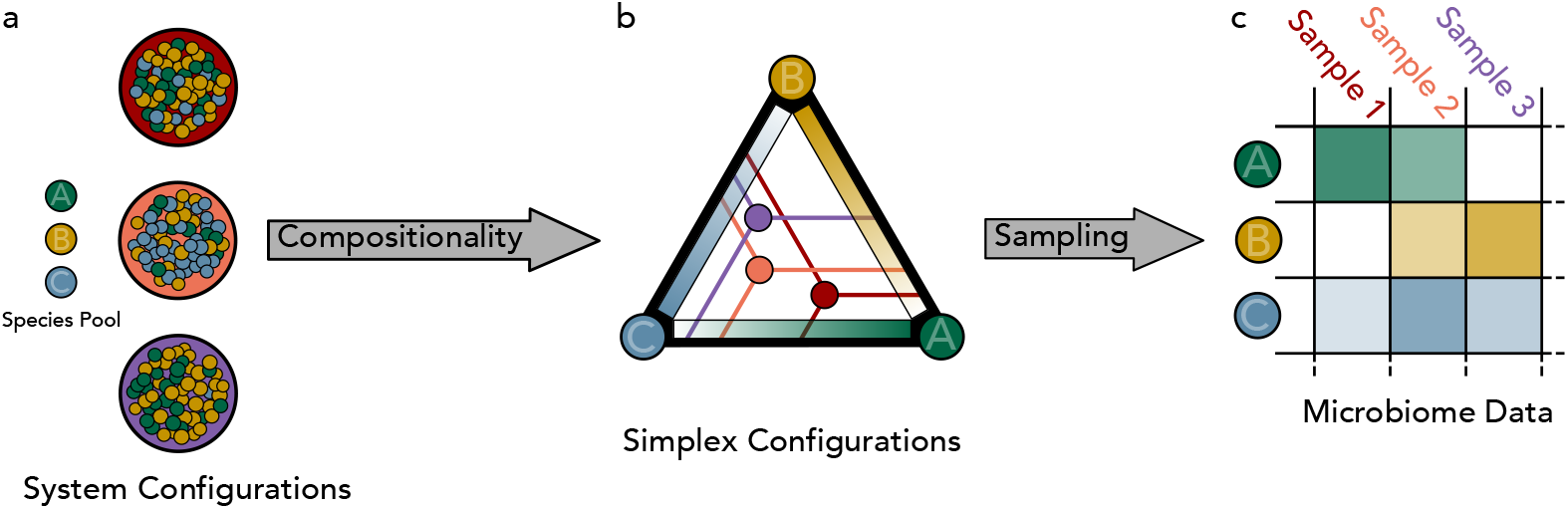
Schematic representation of the proposed theoretical framework to generate synthetic taxonomic data tables. Panel a: colored large circles indicate different samples of microbial communities. Each small ball inside the circle represents an individual of a given species (A,B,C). We normalize the species abundances *n*_*i*_ to densities *v*_*i*_ 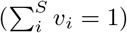. In Panel b we show the densities phase space for the case of three species and three samples, which can be represented by a simplex. Each point inside the simplex now represents one of three samples, while each vertex represents a configuration in which only the corresponding species is present in the samples. The relative abundance contribution of each species can be represented by a gradient. For a given sample, in order to obtain the relative abundance of a species, one has to project parallel to the simplex sides (as denoted by the lines). Panel c: we eventually introduce sampling, and generate the taxonomic tables through a multinomial distribution with the densities *v*_*i*_, and the number of individuals as those observed in the empirical data. Species with low relative abundance may not be sampled, such as species B in sample 1.

We will compare the results obtained for both MD and MSSD models and also for the compositional-only-on-average stochastic logistic model originally proposed by Grilli [13] (SLG), i.e., 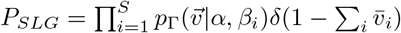. Since the corresponding joint species abundance distribution does not have a sum-to-one hard constraint, the appropriate sampling distribution will be the Poisson, whith which we conovolve the model, obtaining the Poisson Stochastic Logistic Growth model (PSLG). In this model, the ingredient that ensures (on average) compositionality is the fact that mean abundances are constrained to sum up to one. We note that, from a statistical mechanics perspective, *P*_*ζ*=2_ and *P*_*SLG*_ are, respectively, the microcanonical and canonical formulations of the same model. Similarly to the MD, the parameters required to fully specify the model are *µ*, λ and *A*.

Operatively, in order to sample from the three null models, we implement the following procedure: 1) Depending on the model, fit *µ*, λ, *A* from the data (for a detailed discussin abouot fitting procedures, see section 4 of the Supplementary Materials). 2) Extract S average abundances from *P*_MAD_(*µ*, λ); 3) Using Eqs. (6) and the estimate of the coefficient *A* from the TL, generate 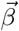 and 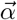 (whose dimensions depend on *ζ*, see equation (6)); 4) Sample the relative species abundances *v*_*i*_, *i* = 1, ..*S*_*R*_ for each of the *r* = 1 … *R* samples we aim to generate using Eq. (7) or Eq. (8); 5) For each sample, with the appropriate sampling distribution, generate species abundances counts using relative abundances and *v*_*i*_ and *N*_*r*_; 5) Remove all species that have relative abundance below a given threshold *κ*, i.e., *v*_*i*_ < *κ*. We will apply three different cut-off *κ*_*l*_ = 4.5 *×* 10^*−*7^ (low), *κ*_*m*_ = 9 *×* 10^*−*6^ (medium), *κ*_*h*_ = 1.8 *×* 10^*−*4^ (high). The three cut-off values were obtained considering how the diversity of the dataset changes with *κ* (for a derivation of these values see Supplementary Materials, Figure 5).

Equipped by these null models, our aim is to investigate the statistical properties and macroecological patterns of two different groups of human gut microbial communities. The first is a cohort of healthy (H) individuals, while the second is of individuals affected by gastrointestinal tract diseases, which we will generally refer to as unhealthy (U).

### Microbiome Metadata Selection and Analysis

We have selected gut microbiome data from three studies [4, 10, 25], where both sequencing data and sample metadata are available for controls and three gastrointestinal tract diseases: Crohn’s Disease, Ulcerative Colitis and Inflammatory Bowels Syndromes. In the following, we will refer to controls as the H group and all samples from pathological conditions as U group. We have filtered so to have a homogeneous and not biased dataset (see section 5 of the Supplementary Materials for details). In general, we have selected the patients less affected (at least at the time of the study) according to medical treatments, to limit the impact of different drug treatments on gut microbiome. After this filtering procedure, we ended up with *R*_*H*_ = 91 shotgun metagenomic samples from healthy control individuals and *R*_*U*_ = 202 samples of dysbiotic microbiome.

We have implemented a computational pipeline to process throughput sequencing metagenomic data following best practices [35], such as quality filtering (remove reads with *Q* < 20) and human DNA decontamination with the NCBI (GRCH38) human genome assembly (GRCH38) ^1^.

The metagenomic taxonomic profiling tool we have adopted in our analysis is the *Kaiju* classifier [36], which converts metagenomic reads in all possible open reading frames and searches for the best match in a protein database. The advantage of this procedure is that, thanks to the degeneracy of genetic code, it is robust to random mutations along the genome and, as such, to evolutionary divergences between the dataset and the reference catalog of genomes. As a reference species catalog, we have used RefSeq [37], which contains protein sequences from complete archaeal and bacterial genomes. Metagenomic samples were profiled on October 13th 2020. Eventually, we have classified (on average across samples) 39% of reads in H samples and 37% U, at the species level. From these we build two data-tables, one for each of the two classes (H and U), having as rows the different species (S) and as columns (R) the samples.

Each {*i, j*} entry gives the corresponding relative species abundances *v*_*i*_ of the sample *j*, so that 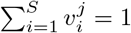. Relative abundances are obtained by dividing the number of reads assigned to a given species by the total number of reads recognized at the species level for that sample. To implement the relative abundance threshold, we set to zero all species abundances that are smaller than the relative abundance cut off *κ*, i.e. 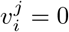 if 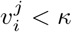.

## Results

### Mean Abundance Distribution and Taylor’s Law

We find that an emergent *MAD* is observed in both H and U datasets. However, we observe a dependence of the shape of the distribution depending on the relative abundance cut-off *κ*. In particular, for *κ* < 10^*−*5^ the MAD displays a *Log-Laplace* shape, i.e. P 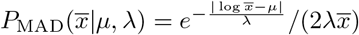, while for *κ* > 10^*−*5^ the MAD is a Log-Normal distribution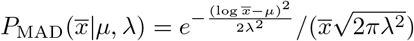, the same found for OTU 16s data [13]. These distributions indicate high heterogeneity in mean abundances having both a heavy tail. In the y-axis of Figure 2 we show the Byesian Iformation Criterion (*BIC*) ratio: if it is greater than one, it indicates that the Laplace distribution is a better fit than the Log-Normal, while if *BIC* <1, then the opposite is true. We thus obtain the values of *µ* and λ for three different thresholds of *κ*. As also clear from Fig. 2, the Laplace distribution is usually a better description for the MAD, except for a large threshold where the MAD clearly displays a Log-Normal shape (and compatibly with the OTU case [13]). As explained in section 4 of the Supplementary Materials, the MSSD model is insensitive to the value of *µ* to generate synthetic table, while the value of λ is inferred from the data and we find λ_*H*_ = 1.407 *±* 0.005 and λ_*U*_ = 1.413 *±* 0.007 (uncertainty of the fit evaluated with bootstrap procedure) as the best fit for the mean species abundances obtained without threshold.

**Fig 2.**
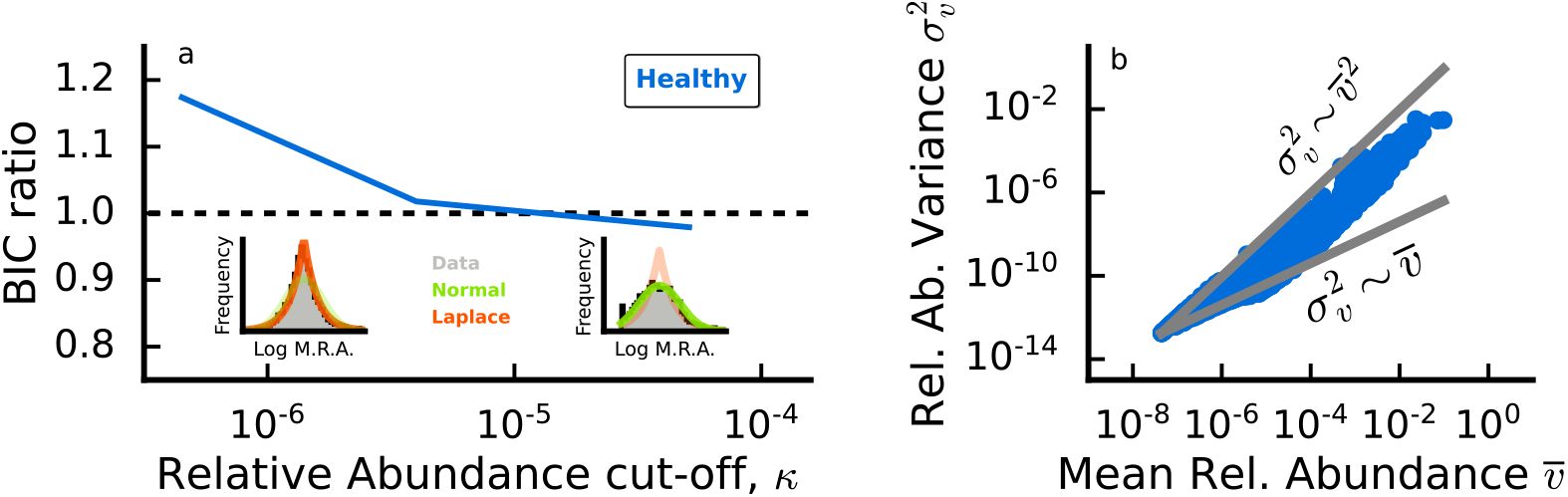
Panel a: *Mean Abundance Distribution* shape displays a dependence on the relative abundance cut-off *κ*. BIC ratio curve for healthy and disease-related data collapse onto the same line. Panel b: Taylor’s Law holds in empirical human gut communities, both in health and in disease. For rare species, it is difficult to discriminate between Poisson-like and Taylor-like scaling, due to the fact that rare species are present only in a few samples. For simplicity, we report the scatter plot only in the healthy case. A brief discussion on the fit procedure can be found in the Supplementary Materials, section 4.

Regarding the TL, we find threshold dependence of the amplitude *A*, while the exponent *ζ* ≈ 2 is remarkably robust. In particular, for each value of *κ* we compare the *R*^2^-score ratio of the best fit power law with the one with fixed *ζ* = 2 finding 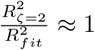 suggesting a negligible discrepancy between the two models for this scaling relation.

Thus, in what follows we will assume *ζ*_*Data*_ = 2.

### Emergent Ecological Patterns in healthy and unhealthy microbiomes

In this section, we investigate macro-ecological emergent patterns in gut microbiomes, and test whether our null model can describe them, and if there are any statistically significant deviations in such patterns between H and U samples.

We will focus on the following ecological patterns of the gut microbiomes: 1) *α* and *γ* diversity [38], defined as the number of different species in each local community (i.e., samples) and H and U meta-communities (i.e. union of all H/U samples), respectively; 2) The abundance-occupancy distribution, describing the probability for a species with mean relative abundance 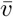 is found in a fraction 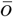 of the total number of samples within a meta-community; 3) The species abundance distribution (SAD) of the H and U meta-communities; 4) The relation between the number of species observed in a local community normalized to the meta-community one (*α*/*γ*-diversity) and its sequencing depth (i.e. the metagenomic version of species-area relationship).

To compare a given ecological pattern obtained from the data and the corresponding one produced by the null model, we first set the scaling exponent *ζ* (*ζ* = 2 for MSSD and SLG, *ζ* = 1 for MD). Second, we fit the parameters *µ*, λ, A (in the case of MSSD we do not fit µ due to model symmetries) from the data with no cut-off (*κ* = 0). Then we generate 500 realizations of the two meta-communities (H and U) with the same number of reads (*N*), of species (*S* = *γ*), and of samples *R* as found in the data.

Eventually, we consider the three relative abundance cut-off *κ* both in the empirical and simulated data (i.e. all species with *v*_*i*_ < *κ* is set to zero) and for each pattern we calculate a *R*^2^-like score (see Supplementary Material, section 3). The final *R*^2^-score we assign to the model is the average over the instances of the model.

The first patterns we consider are the *γ* and *α* diversity. The values of such quantities strongly depend on *κ*, but reveal a persistent regularity that is independent of it. Indeed, the overall (*γ*) diversity of species is found in the unhealthy meta-community is larger than the one found in healthy microbiomes (*γ*_*H*_ < *γ*_*U*_). On the other hand, we have that the *average* local diversity 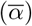 found for H samples is larger compared to one for U samples, i.e., 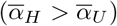 (see Figure 3)

**Fig 3.**
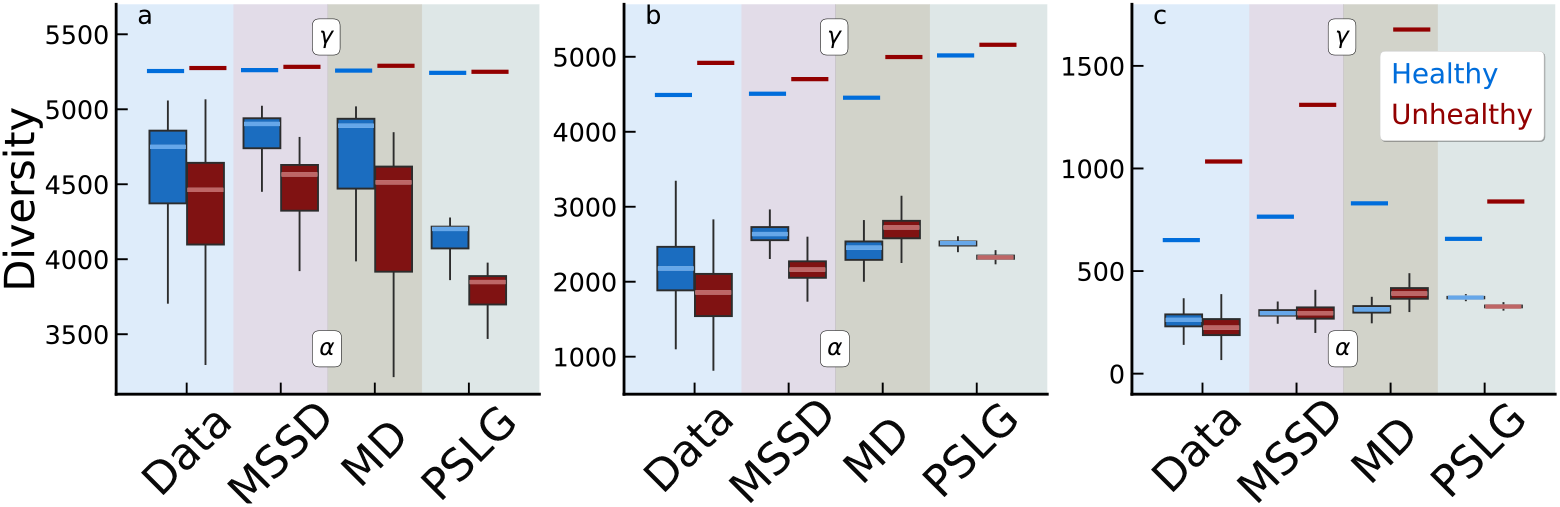
Box-whiskers plots describe average local (α) diversity, while horizontal bars indicate the corresponding metacommunity (*γ*) diversity. In healthy gut microbiomes (in blue) we find higher average *α* and lower *γ* diversity than unhealthy ones (in red). The three panels represent different threshold relative abundance cut-off *κ*: a) low (*κ*_*l*_ = 4.5 × 10^*−*7^); b) medium (*κ*_*m*_ = 9 × 10^*−*6^); c) high (*κ*_*h*_ = 1.8 × 10^*−*4^). In each panel, we compare the diversity of the empirical H and U diversity with respect to the one generated by three different null models: MSSD, SLG, and MD. MSSD is found the best model, especially for low and medium *κ*.

The second pattern we consider is the SAD [26], which describes, in a given sample the probability of observing species with a given abundance. In agreement with previous results obtained with 16s OTU data [13, 39] and also shotgun data [40], we find that the SAD displays a heavy tail, that is compatible with small and medium cut-off with power-law distributions with exponents around 1.7 (see Figure 7 of the Supplementary Materials for more details). For large thresholds, the SAD is more compatible with a Log-Normal distribution. No significant differences are observed between the H and U individuals (see Figure 4a). Moreover, all the models (PSLM, MSSD, and MD) generate SADs that are compatible with the empirical ones. These results confirm [41] that SADs are not informative patterns of the underlying ecological mechanisms driving species abundances.

**Fig 4.**
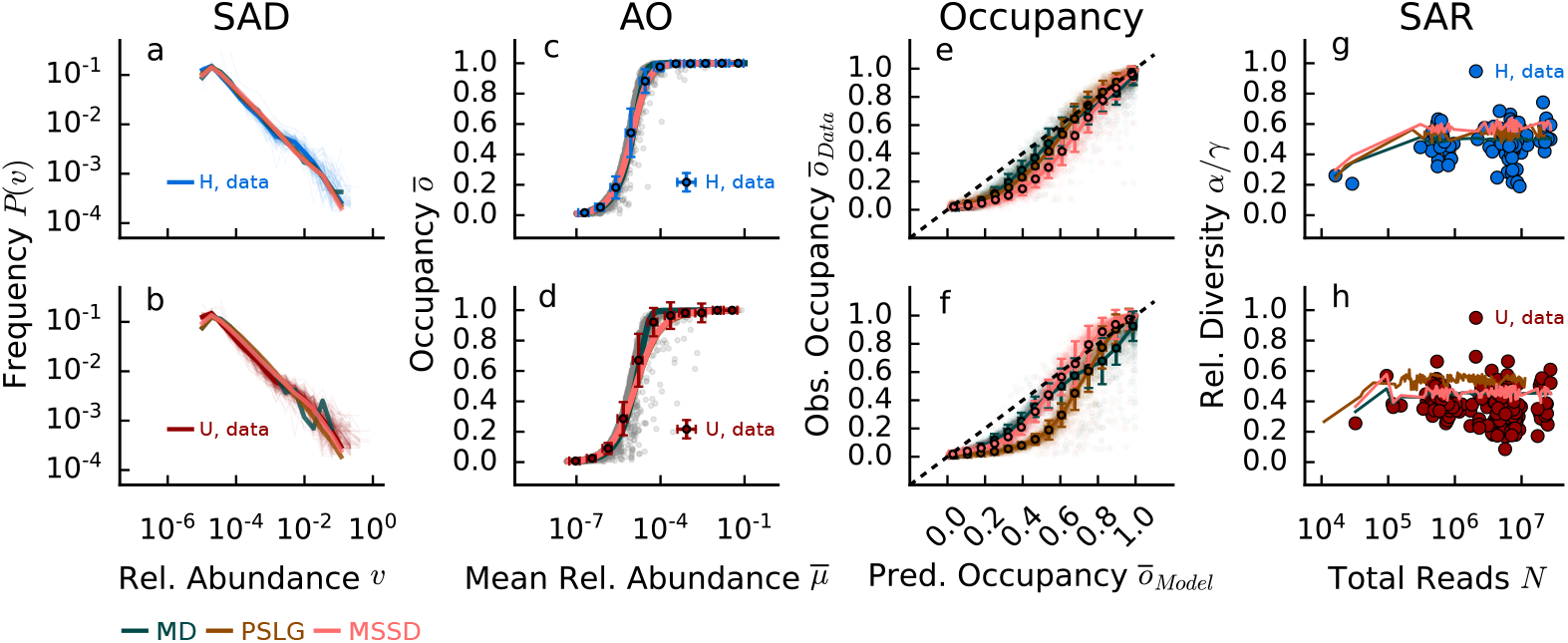
Comparison of emergent empirical ecological patterns in healthy (H, top panels) and unhealthy (bottom panels) microbiomes and for MSSD, SLG and MD models. Panels a-b) Species Abundance distribution (SAD); c-d) Species abundances occupancy curve (AO). Grey shaded points refer to single species mean relative abundance and occupancy; e-f) Empirical vs. predicted occupancy curves (O). Shaded points refer to the predicted/observed occupancy of single species; g-h) Species Area Relationship curves. These patterns can be investigated for different cutoffs (see Supplementary Materials, Figures 1-4), here we show the average threshold cut-off *κ*_*m*_ = 9 *×* 10^*−*6^.

**Fig 5.**
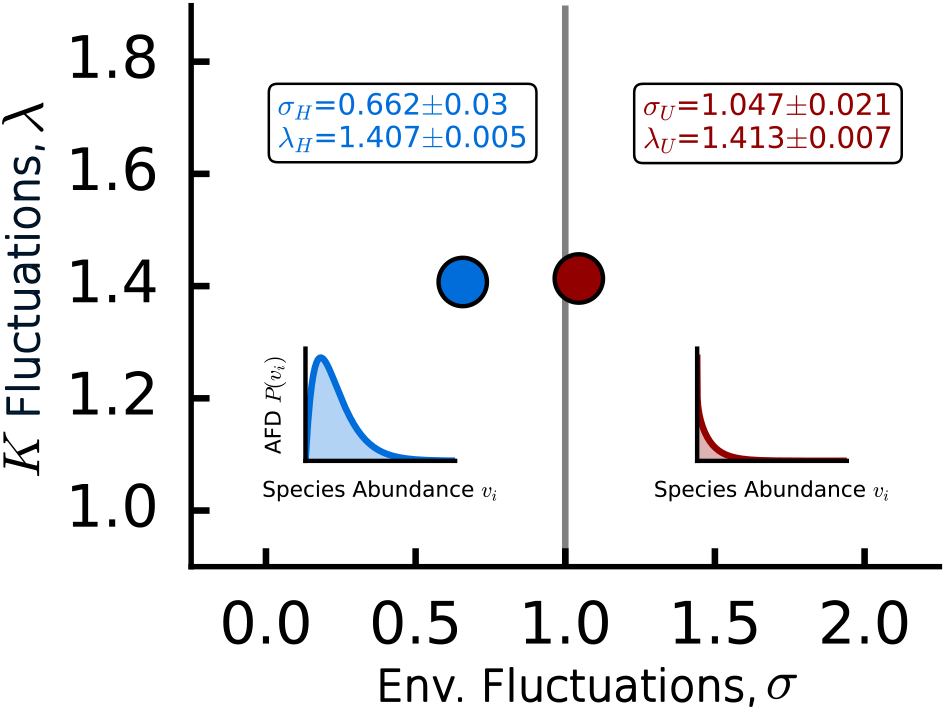
Taylor’s Law and Mean abundance distribution parameters are informative of the underlying stochastic logistic model. The values obtained from the data suggest that healthy and unhealthy species abundance distribution follow different qualitative behaviors, with the unhealthy case being prone to more extinction. Average and standard deviation estimates of the parameters have been obtained through a bootstrap procedure.

We then investigate the relation between the log-mean relative abundance of a species and its occupancy, which we refer to as abundance-occupancy (AO) curve. We have already introduced the average relative abundance of species *i* as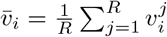, while we define the *occupancy* of species *i* as 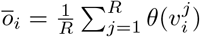, where *θ* is the Heaviside Theta, which converts relative abundance data into presence/absence ones.

The relation between 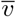 and 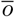, as shown in Figure 4b, describes how likely it is for a species, given its average relative abundance to be sampled in a realization of the system (also known as intensity-sparsity relation [14]). When the low/medium value of *κ* is set, the curve suddenly saturates, suggesting that a vast proportion of the available *S* = *γ* species is expected to be sampled. In this scenario, the community is dominated by rare species, and thus almost all sampled individuals belong to different species, thus saturating the diversity very fast. As *κ* increases we have fewer and fewer species that are rare in relative abundance but, at the same time are harder to sample. As Figure 4c, d shows, this behaviour is shared by all models, and thus is also not discriminative of underlying ecological processes. However, for low (high) *κ* the MD typically underestimates (overestimates) the occupancy of species (not shown here, see Supplementary Materials, Figure 1).

We can also compare directly the occupancy curves obtained from presence-absence data, and those predicted by the models (as shown in Figure 4e-f). Interestingly, contrarily from the previous patterns, in this case, there are differences in how well the models describe the data. In particular, for H samples, the PSLG model outperforms the MSSD one, while on the contrary MSSD better predicts the pattern of U samples. Such results and goodness of fit for the MD model are not robust for different thresholds (see Table 1 and Supplementary Materials, Figure 1).

**Table 1.** *R*^2^ scores for the considered macroecological patterns and for three different thresholds: low *κ*_*l*_, medium *κ*_*m*_ and high *κ*_*h*_

|  |  | AO Curve |  |  | Species Occupancy |  |  | SAR Curve |  |  |
| --- | --- | --- | --- | --- | --- | --- | --- | --- | --- | --- |
| | | $\kappa_l$ | $\kappa_m$ | $\kappa_h$ | $\kappa_l$ | $\kappa_m$ | $\kappa_h$ | $\kappa_l$ | $\kappa_m$ | $\kappa_h$ |
| H | MSSD | 0.997 | 0.996 | 0.983 | 0.850 | 0.823 | 0.830 | 0.819 | 0.695 | -15.131 |
|  | PSLG | 0.998 | 0.995 | 0.983 | 0.908 | 0.912 | 0.811 | 0.984 | 0.585 | -22.655 |
|  | MD | 0.937 | 0.995 | 0.944 | 0.516 | 0.925 | 0.337 | 0.776 | 0.005 | -59.763 |
| U | MSSD | 0.992 | 0.982 | 0.987 | 0.885 | 0.861 | 0.776 | 0.976 | 0.840 | -0.779 |
|  | PSLG | 0.987 | 0.982 | 0.984 | 0.911 | 0.571 | 0.569 | 0.988 | 0.522 | -1.244 |
|  | MD | 0.902 | 0.993 | 0.864 | 0.562 | 0.872 | 0.485 | 0.874 | 0.726 | -1.302 |

We finally consider the ”metagenomic” version of the species-area-relation (SAR) [42], i.e. how the diversity increases with increasing sampled area. Here the area is substituted by the total number of classified reads. Therefore, we consider increasing the number of reads combining the samples of each group and calculate the normalized diversity as the number of unique species in the aggregated community divided by *γ* diversity. Although all models on average slightly overestimate the overall diversity, in H samples they all perform similarly, while for U samples MSSD and MMD models (that can generate the same number of total reads as found in the data) are better than the SLG model.

Table 1 summarizes all the results and the comparison between the goodness of fit (measured as explained in section of the three models for the presented emergent empirical ecological patterns of gut microbiomes in both health and disease states.

Finally, by fitting the MSSD, we can gain insight into possible differences in species population dynamics within microbiomes from healthy and diseased individuals. As reported in Figure 5, by fitting the MAD we find λ for both H and U cohorts. We can interpret it as the fluctuation scale of the carrying capacities. This turns out to be statistically indistinguishable between the H and U cohorts. However, the most relevant difference comes from the value we infer for *σ*, the width of the environmental fluctuations. The PSLG predicts that the abundance fluctuations distribution has a polynomial part with an exponent greater than zero if *σ* > 1 (as observed in the unhealthy cohort) and less than zero if *σ* < 1 (as observed in the healthy cohort).

## Discussion and Conclusion

By inferring the parameters of the model best describing these patterns, we have obtained ecological insights into dysbiosis that would not be directly accessible from the data. In particular, we have found that while the intrinsic population dynamics are similar in the H and U cohorts (they have a very similar carrying capacity determined by λ), fluctuations in growth rates due to extrinsic environmental factors (given by σ) are much stronger in the U microbiomes. This is reflected in the abundance fluctuations distribution shifting from a modal distribution in H samples to a power-law one (with exponential cut-off) in U microbiomes, that is, in dysbiosis the probability for a species to be very rare is higher.

This result allows us to explain why in the H cohorts we observe a higher *α* but a lower *γ* diversity: the species in the U microbiomes experiencing higher fluctuations are more prone to extinction locally, but also are subject to higher turnover (thus increasing the global diversity of the group). Thus, we believe that for dysbiois the celebrated Anna Karenina principle ^2^ also holds: ’All healthy gut microbiomes are alike; each unhealthy gut microbiome is unhealthy in its own way”. In fact, there are convergent observations suggesting that dysbiosis can be attributed to host-specific factors [43, 44]. We also have implemented a stratification analysis (see Supplementary Information, Section 7), where we investigated whether the distinct diversity patterns for healthy and unhealthy microbiomes also hold if we consider only specific gastrointestinal diseases.

We have observed that while Chron disease (CD) and ulcerative colitis (UC) exhibit trends consistent with our overall findings (lower average α diversity and higher *γ* diversity in U patients), inflammatory bowel syndrome (IBS) presents less systematic tendencies. This distinction is noteworthy given the clinical challenges associated with the diagnosis of IBS. We have also performed a stratified inference of the environmental noise (*σ*) and carrying capacity heterogeneity (λ) parameters, finding results compatible with the previous ones. For the different cohorts, there is no substantial difference in λ; On the other hand, we have found that CD and UC are associated with an environmental noise strength close to one and larger than healthy microbiomes, supporting our interpretation discussed above. Again, IBS has a behavior more similar to that of healthy individuals.

We have also shown that microbiome species abundance data exhibit a Taylor law with a *ζ* value of 2. Interestingly, the MD model, by its design, does not satisfy this fundamental constraint. However, as shown in Table 1, the MD model is capable of generating AO curves and SADs that are compatible with the data at any threshold. Similarly, it can accurately reproduce Occupancy curves at a medium threshold and SAR curves at a low threshold. This finding highlights that not all patterns and thresholds are equally informative. Some are more effective than others in differentiating between models and underlying ecological processes. However, AO curves have previously been used to test specific underlying ecological theory [45]. On the contrary, our results suggest that the shape of the AO curve is simply the result of two main ingredients: heterogeneous population averages and random sampling. Similarly, all patterns at low thresholds are dominated by a vast number of rare species (that in shotgun data are probably false positives [15]).

Something similar occurs for the SAD, where all models are practically indistinguishable from one another. Indeed, the fact that different models (i.e., processes) can lead to very similar SAD patterns has long been known in theoretical ecology [26, 46].

For the SAR curves there is a very strong impact of the thresholds on the models goodness of fit. At low thresholds, all models performed well, thus not providing any discriminatory power for the right choice of the model. However, at the high threshold, there was a significant drop in the goodness of fit for all models. In fact, due to removal of all the rare species, the SAR loses its characteristic shape, and thus it is not useful for models comparison. At intermediate threshold, the MSSD model performed the best, although it is important to note that the *R*^2^ value is not particularly high. In this case, we also have found that all models fit better to U samples than to H samples. This effect is probably because in U samples the *γ* diversity is higher, and we have many more rare species, thus increasing the slope of the SAR (that is general overestimated by the models).

Upon closer examination of the Species Occupancy patterns in Table 1, notable differences emerged between healthy and unhealthy samples at the medium threshold. In the H samples, the SLG model showed the highest goodness of fit, closely followed by the MSSD model. On the contrary, the unhealthy samples showed a different pattern. The SLG model, which performed strongly in the healthy samples, showed a marked decrease in its goodness of fit, indicating potential challenges in capturing the complexities of species occupancy in unhealthy systems at this threshold. The MSSD model also showed a reduction in performance, but remained relatively more consistent compared to the SLG model. The performance of the MD model for both H and U samples was extremely variable, depending on the cut-off *κ*. For medium threshold, the fit - although not as good as the one of the MSSD and SLG models - had a relatively high *R*^2^, while for low and high thresholds, it decreased markedly.

In our study, we thus have found that, contrary to AO curves and SADs, species occupancy and diversity curves provide key insights into the performance of various models. Models incorporating Taylor’s law with *ζ* = 2 (SLG and MSSD) offer a better explanation of the data at the medium threshold (which has proven to be the most informative cut-off for false positive). This suggests that large species fluctuations, as dictated by Taylor’s law with a scaling exponent of *ζ* = 2, are important for accurately predicting presence/absence patterns and species diversity in empirical datasets. We also have found that models performing well in healthy communities may not necessarily do so in unhealthy ones, and vice versa. This insight is crucial for ecological modeling and could guide future research in developing or choosing models that are tailored to the specific conditions of the ecological systems being studied. Furthermore, we have found that the SLG model, although effectively similar to the MSSD in many respects, underestimates species occupancy (see Figure 4f) and overstimates species diversity (see Figure 4g). The reason is that SLG can generate ecological communities with a number of individuals that is only on average as the one of the corresponding sampled data, while MSSD implements strict compositionality of the data.

All in all, we suggest that although compositionality and sampling strongly obscure ecological signals making the majority of empirical patterns qualitatively similar, there are indeed quantitative ecological differences between microbial communities of the gut microbiome in health and disease. In particular, by considering only a few relevant patterns like Taylor’s law and species occupancy and by using interpretable analytical models that also include environmental noise, we may propose an interpretation of the observed differences in the taxonomic data, eventually shedding light on underlying ecological processes characterizing informative emergent patterns, such as the specific trend of *α* and *γ* diversity in both H and U cohorts. We thus conclude that dysbiosis is characterized by stronger turnover than healthy microbiomes, which is due to larger environmental fluctuations and also reflected in marked heterogeneities of species’ carrying capacities.

## Supporting information

**S1 - List of abbreviations**

**S2 - Supplementary figures**

**S3 - Model Testing**

**S4 - Supplementary Methods**

**S5 - Data Selection, Processing and Analysi**.

**S6 - Metagenomic Pipeline Implementation**

**S7 - Stratification Analysis**

## Acknowledgments

S.S. acknowledges Iniziativa PNC0000002-DARE - Digital Lifelong Prevention. A.M. also acknowledges the support of the NBFC to the University of Padova, funded by the Italian Ministry of University and Research, PNRR, Missione 4, Componente 2, “Dalla ricerca all’impresa”, Investimento 1.4, Project CN00000033. A.R. acknowledges funding from Fondazione Cassa di Risparmio di Padova e Rovigo (IT) through its grant 55722.

## Footnotes

1 https://www.ncbi.nlm.nih.gov/datasets/genome/GCF_000001405.26/

1 ”All happy families are alike; each unhappy family is unhappy in its own way” (Lev Tolstoj, Anna Karenina, 1877)

